# SN-FPN: Self-attention Nested Feature Pyramid Network for Digital Pathology Image Segmentation

**DOI:** 10.1101/2024.04.30.591883

**Authors:** Sanghoon Lee, Kazi A. Islam, Sai C. Koganti, Varshini Yaganti, Sai R. S. Mamillapalli, Hannah Vitalos, Drew F. Williamson

**Author notes:** Corresponding author: Sanghoon Lee.

## Abstract

Digital pathology has played a key role in replacing glass slides with digital images, enhancing various pathology workflows. Whole slide images are digitized pathological images improving the capabilities of digital pathology and contributing to the overall turnaround time for diagnoses. The digitized images have been successfully integrated with artificial intelligence algorithms assisting pathologists in many tasks, but there are still demands to develop a new algorithm for a better diagnosis process. In this paper, we propose a new deep convolutional neural network model integrating a feature pyramid network with a self-attention mechanism in three pathways: encoder, decoder, and self-attention nested for providing accurate tumor region segmentation on whole slide images. The encoder pathway adopts ResNet50 architecture for the bottom-up network. The decoder pathway adopts the feature pyramid network for the top-down network. The self-attention nested pathway forms the attention map represented by the distribution of attention scores focusing on localizing tumor regions and avoiding irrelevant information. The results of our experiment show that the proposed model outperforms the state-of-the-art deep convolutional neural network models in terms of tumor and stromal region segmentation. Moreover, various encoder networks were equipped with the proposed model and compared with each other. The results indicate that the ResNet series using the proposed model outperforms other encoder networks.

## I. INTRODUCTION

**T**RADITIONAL pathology examines tissue samples using a microscope by a pathologist to diagnose diseases such as cancer, infectious diseases, and hematologic disorders, determining the characteristics of their development of abnormal cells [1], [2]. However, these examinations have been limited to manual examination of slides, subjective interpretation, and not easily shareable slides, causing time-consuming, subjective, and collaborative challenges. Digital pathology is a branch of pathology assisting traditional pathology by providing high-resolution digital slide images through digital slide scanners and image analysis tools quantifying and interpreting pathology data [3], [4]. Adopting digital pathology can enable pathologists to improve their workflow efficiency, accessibility, and quantitative analysis, advancing biomedical knowledge and research.

Whole slide images (WSIs) are digital slides representing entire pathology slides with high resolutions approximately 1.5 gigabytes per slide allowing detailed examination of tissue structures [5]. WSIs are typically obtained by a process of digital slide scanning converting glass pathology slides stained using various histological stains (i.e., hematoxylin and eosin (H&E)) into digital images at high magnification [6]. Since WSIs are the digitized representation of entire pathology slides, they enable pathologists to explore the detailed information of tissue samples across the entire image, enhancing the examination process of pathology images by collaborating with other pathologists and integrating with image analysis techniques [7]. As WSIs become a key component of digital pathology, it is necessary to develop a new methodology to enhance the whole slide image analysis.

The analysis of WSIs to facilitate research in digital pathol-ogy has been developed through advanced image analysis techniques such as object detection, image classification, and image segmentation [8]. Object detection techniques in digital pathology have been utilized for automatically detecting regions of interest such as tumors, immune cells, and nuclei [9], [10]. A typical object detection process includes data preprocessing, training a model, validating a model, indicating the spatial location (i.e., bounding box), and post-processing. Image classification techniques in digital pathology have been utilized for automatically categorizing or classifying entire images into a set of labels [11], [12]. Image classification techniques are very similar to object detection techniques but assign a single label to the entire patch or image. Image segmentation techniques have been used in digital pathology to delineate different regions based on visual context. While object detection and image classification aim to obtain the object or class label, image segmentation is interested in partitioning an image pixel into meaningful or homogeneous regions [13], [14]. In this paper, we focus on image segmentation techniques in digital pathology.

Traditional image segmentation approaches have mainly utilized hand-engineered models grouping pixels with similar properties or identifying boundaries between regions [15], [16]. These models highly rely on handcrafted features such as edges, corners, and even entropy, which are computationally simple but struggle with scalability and effectiveness on complex features. To remedy these issues on hand-engineered models, deep learning-based models have been introduced for capturing complex representations and contextual information from raw data aiming at accurate segmentation of regions [17], [18].

Deep-learning models, particularly convolutional neural networks (CNNs), have significantly contributed to image segmentation [19], [20]. The convolutional layers in CNNs can enable the deep-learning model to identify local patterns of spatial information in specific regions, the down-sampling layers interspersed between the convolutional layers can down-sample the spatial dimension of the prior map, reducing computational complexity and building spatial hierarchies of features, and the up-sampling layers recover the spatial resolution equal to the input image. CNNs automatically excel at learning hierarchical and spatial features from input images through the combination of convolutional, down-sampling, and up-sampling layers.

Feature map generation at different scales using CNNs has been a widely accepted approach for deep learning-guided applications such as object detection, image classification, and image segmentation [21]–[23]. Feature maps obtained from multi-resolution representations of image pyramids can be independently used for image prediction or the feature maps can be combined with the skip connections producing a single-level feature map through a top-down architecture called Feature Pyramid Network (FPN) [24]. However, using feature maps for every prediction requires a large amount of computation and the top-down model with the skip connections remains to improve the effectiveness of the applications.

In this paper, we propose a feature pyramid network combined with a concept of a self-attention model equipped over the last state of the encoder creating an attention map for the first state of the decoder. We named it the Self-attention Nested Feature Pyramid Network (SN-FPN) for digital pathology image segmentation. SN-FPN takes advantage of self-attention mechanisms handling irregular shapes or contours of image regions and capturing contextual relevance and spatial relationships between pixels in an image. The differences between the proposed model with other models are briefly shown in Fig 1. Features are merely extracted from each image pyramid for predictions (left). Features are extracted by using the feature pyramid network through the bottom-up and top-down pathways; the features on the top-down pathway are merged with the corresponding features on the bottom-up pathway (middle). The proposed self-attention mechanism is nested between the bottom-up and the top-down pathway (right). We describe the details of the proposed SN-FPN model in Section III.

**FIGURE 1.**
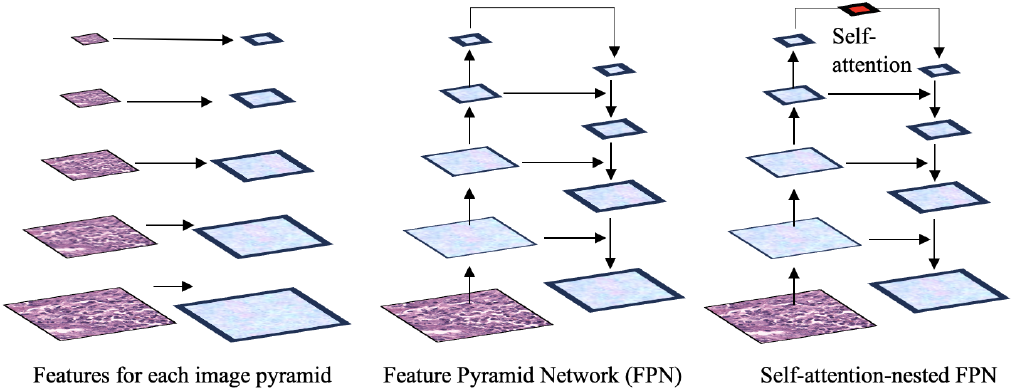
Comparision of Image Pyramid, Feature Pyramid Network, and the Proposed Self-attention Nested Feature Pyramid Network.

## II. RELATED WORKS

FPN is a multi-resolution feature pyramid network that has been widely studied in various types of models. Mask Region-based Convolutional Neural Network (R-CNN) adopted the concept of multi-scale feature maps of FPN to improve its performance in object detection tasks by capturing object information at different scales [25]. RetinalNet is an object detection model that aims to solve the class imbalance problem by assigning different weights on different scale objects using the Focal Loss [26]. To address the problem, RetinalNet employed FPN as a backbone architecture to enable the top-down architecture with lateral connections thereby capturing multi-resolution semantic information. Cascade R-CNN is an extended version of Faster-RCNN, using a series of detection branches in a cascaded manner refining the results of the prior stage [27]. Both FPN and Cascade R-CNN are similar in terms of multiple stages but Cascade R-CNN follows a cascaded structure. Neural Architecture Search for FPN (NAS-FPN) is an extended version of FPN combined with the concept of Neural Architecture Search. NAS-FPN aims to automatically design feature pyramid architecture through the optimization of the FPN architecture [28]. High-Resolution Network (HRNet) is a multi-scale convolutional neural network that maintains high-resolution images through parallel multi-resolution convolutions and repeated-resolution fusions [29]. Path Aggregation Network (PANet) is an instance segmentation deep convolutional neural network aiming to improve the feature pyramid network by taking accurate localization information at low levels [30]. However, these related works mainly focus on object detection or pose estimation and do not handle attention mechanisms.

Attention Aggregation-based Feature Pyramid Network (*A*^2^-FPN) utilized the attention mechanism for its network pipeline to improve the multi-scale feature aggregation [31]. Multi-attention Object Detection Model (MA-FPN) took the advantage of attention mechanism by adding pixel feature attention structure through the multi-scale convolution branches [32]. Position Attention Guided Connection Network (PAC-Net) emphasized position attention by capturing salient dependencies with accurate location information for an effective 3D image detection model [33]. Although these methods utilized attention mechanisms and improved FPN, their approaches have been limited to adopting attention at multi-resolution. Moreover, the demonstration of the effectiveness of attention-equipped FPN is also limited for histopathology image segmentation.

## III. SELF-ATTENTION NESTED FPN FOR DIGITAL PATHOLOGY IMAGE SEGMENTATION

In this section, we describe the proposed SN-FPN model for digital pathology image segmentation. First, we will describe the preprocessing steps for whole slide image segmentation. This step will explain how the input source of the SN-FPN can be obtained from whole slide images. Second, we will present the encoder pathway of the SN-FPN. Third, we will explain how the self-attention mechanism can be nested over the output of the encoder pathway. Last, we will show the decoder pathway of the SN-FPN.

### A. WHOLE SLIDE IMAGE SEGMENTATION

Whole slide images are high-resolution digital images of entire tissue cells. Segmenting whole slide images can be done by partitioning the images into regions of interest. Since the typical size of whole slide images is approximately 1.5 gigabytes, these images are commonly divided into smaller, distinct regions to make processing feasible. In this paper, 151 hematoxylin and eosin-stained (H&E) whole slide images provided by the Breast Cancer Semantic Segmentation (BCSS) dataset were used for the input source of the SN-FPN. 2,355 H&E images were cropped from 151 large regions of interest images obtained at 20x magnification [34]. Since these H&E images were created by different laboratories using different scanners, it is necessary to perform a color normalization reducing variations in color and intensity. Color normalization is a process of standardizing image color to avoid irrelevant color variability and is a critical part of whole slide image segmentation. This process ensures that whole slide images are more interpretable when training a deep-learning model, improving the generalization of the model for a new dataset.

Reinhard color normalization was adopted for whole slide image segmentation [35]. First, we converted the RGB image to Ruderman’s LAB color space to compute the mean and standard deviation of the image intensities for each channel. Since the source image is the RGB image, the number of channels is three. Next, the mean and standard deviation obtained in the LAB color space were used for transforming the image color characteristics to the standard color characteristics for normalization. In this step, the LAB color space was scaled to unit variance with zero mean and rescaled and recentered to match the reference image mean and standard deviation [36]. 2,355 H&E images were cropped from 151 large image regions of interest and used as the input source of the SN-FPN. The scaled pixel value 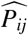 to unit variance for each channel is defined as:

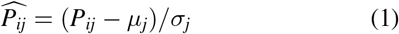

*P*_*ij*_ is the *i*-th pixel value of the *j*-th channel in the LAB color space. *μ* is the mean of the *j*-th channel. *σ* is the standard deviation of the *j*-th channel. The color-normalized image patches were then used as the input source of the encoder pathway. The patches were down-sampled by a factor of 2 for each stage of the encoder pathway and the output of the encoder pathway was used as the input source of the self-attention nested pathway. In the self-attention nested pathway, an attention map was created by using a softmax max function normalizing the attention scores computed by the multiplication of the two feature maps convolved by the filter of 1 by 1 from the output of the encoder pathway. Another feature map convolved by the filter of 1 by 1 was then merged by the attention map multiplied by the convolved feature map. The output of the self-attention nested pathway was used as the input feature map of the decoder pathway. The input feature maps obtained by the attention mechanism were up-sampled by a factor of 2 for each stage of the decoder pathway. These up-sampled feature maps were then merged with the corresponding feature maps with the same resolution convolved by the filter of 1 by 1 in the encoder pathway. These merged feature maps are then used to generate additional feature maps to be merged for the prediction. The details of the whole slide image segmentation using SN-FPN are shown in Fig. 2.

**FIGURE 2.**
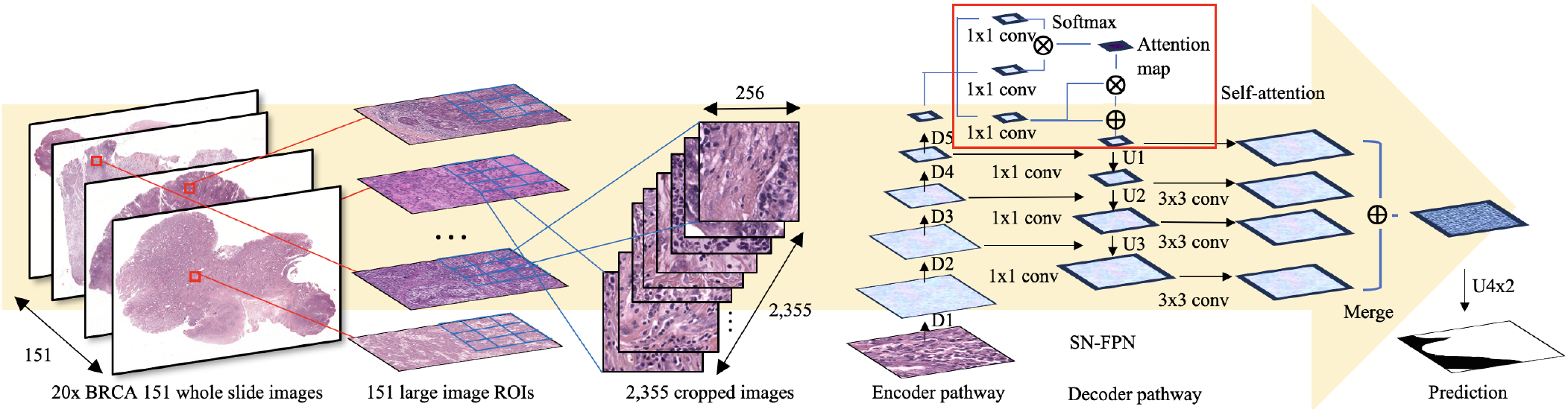
The Overview of the Self-attention Nested Feature Pyramid Network (SN-FPN) for Whole Slide Image Segmentation.

### B. ENCODER PATHWAY

The encoder pathway is the bottom-up network extending from one feature map to another. This bottom-up network performs the feed-forward process consisting of several stages where each stage consists of convolutional layers increasing the number of channels followed by pooling layers which reduce the spatial dimension of the prior feature map, avoiding computational complexity. The output feature map is produced at the last stage of the feed-forward process.

We have adopted ResNet50 [37] for our encoder pathway to produce the output feature map which will later be used as the input source of the self-attention nested pathway. Each stage in the encoder pathway consists of image convolutions followed by pooling or down-sampling by a factor of 2. The number of image convolutions performed in five stages is 1, 3, 4, 6, and 3 respectively. Down-samplings were performed in the four blocks generating 256, 512, 1024, and 2048 feature maps respectively. The details of the encoder pathway are shown in the leftmost blocks in Fig. 3.

**FIGURE 3.**
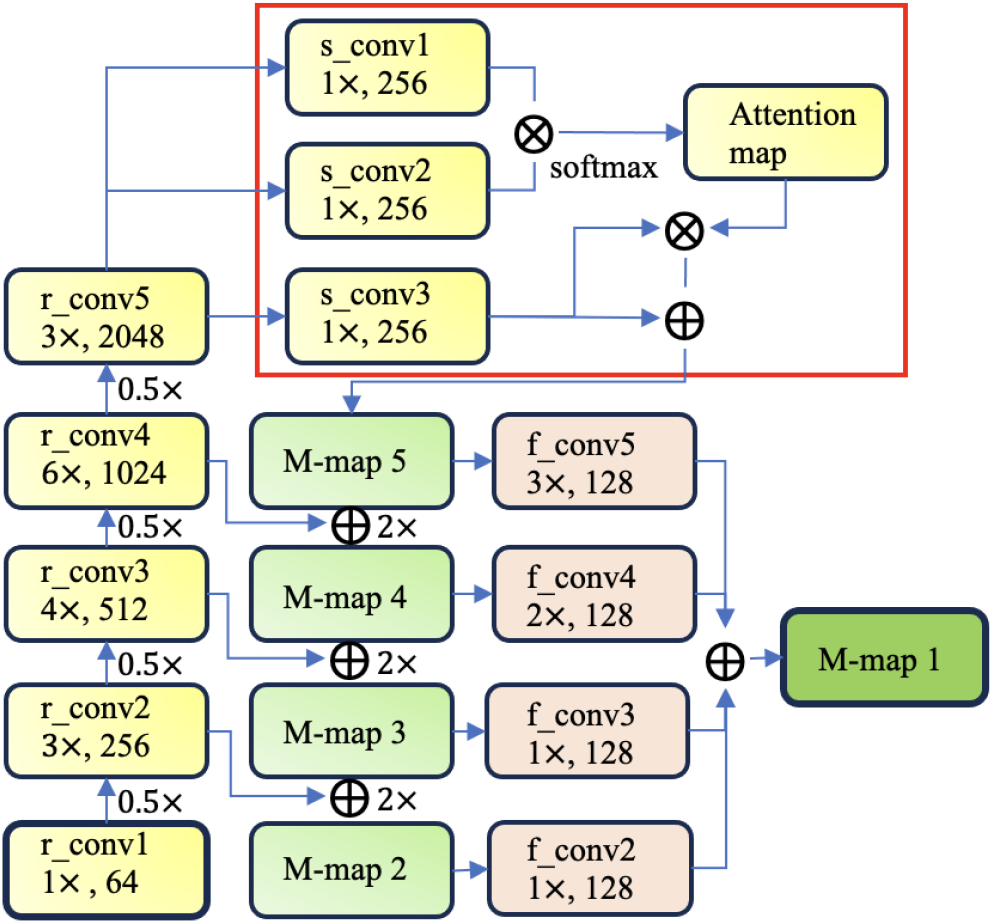
Self-attention Architecture Nested in the Encoder and Decoder Pathways.

### C. DECODER PATHWAY

The decoder pathway is the top-down network extending from one feature map to another. While the encoder pathway down-samples features, the decoder pathway up-samples features mitigating the potential loss of spatial information during down-sampling and facilitating the generation of output images. We have adopted the top-down pathway of the FPN [24], but the output features of the self-attention nested pathway are used as the input features of the decoder pathway. These input features are up-sampled by a factor of 2 and merged with the features of the same size from the encoder pathway. The same-sized features are generated by convolving the filter of size 1 by 1 across the corresponding features from the encoder pathway. The middle blocks in Fig. 3 represent the merged maps with the output of the self-attention nested pathway and corresponding feature maps in the encoder pathway.

While iterating this process until we reach the second feature map, four additional feature maps are generated by convolving the filter of size 3 by 3 across the merged feature maps in the decoder pathway. Because these additional feature maps are intentionally to be the same sizes, they are merged into one feature map for prediction. These additional feature maps are shown as four blocks on the right side of Fig. 3. The rightmost block in Fig. 3 shows the last feature map merged through the four additional feature maps. The last feature map is then up-sampled to be the same resolution as the input image for the prediction.

### D. SELF-ATTENTION NESTED PATHWAY

Self-attention nested pathway is the middle network between the encoder pathway and the decoder pathway. Since the semantic feature values increase as the spatial resolution decreases, we assume that it is necessary to train the model to focus on relevant information rather than on redundant or irrelevant information from the input source thereby leading to accurate and more context-aware prediction results. This process can be done by forming an attention map providing the relatedness of each pixel information in the self-attention nested pathway. The attention map is generated from the attention weights representing the distribution of attention created from the raw attention scores.

To compute the raw attention scores the feature maps in the last block of the encoder pathway were used to derive three feature maps convolved by the filter of size 1 by 1, corresponding to the concepts of query, key, and value in the attention mechanism. The raw attention scores were then computed by matrix multiplication of two feature maps: query and key. After obtaining the raw attention scores, we computed the attention weights by using a softmax function across all feature maps forming an attention map. The attention map was then multiplied by the feature map called value and the results of the multiplication were merged with the value to create the merged feature map used as the input source of the decoder pathway. The merged feature map *F*_*m*_ in the self-attention nested pathway is defined as:

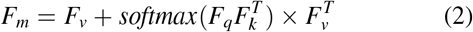

where *F*_*v*_ represents a feature map that plays a role as a value, *F*_*q*_ represents a feature map that plays a role as a query, and *F*_*k*_ represents a feature map that plays a role as a key. The details of the self-attention nested pathway are shown at the top of Fig. 3

## IV. EXPERIMENT RESULTS

In this section, we will perform experiments to verify the effectiveness of the proposed SN-FPN model by comparing it with the baseline of the model and the state-of-the-art models. Moreover, we will explore the effectiveness of the various encoder pathways by adopting deep-learning networks.

### A. H&E IMAGE SEGMENTATION FOR TUMOR REGIONS

#### Dataset

The Breast Cancer Semantic Segmentation (BCSS) dataset is a H&E image dataset that contains more than 20,000 segmentation annotations of tissue regions obtained from breast cancer whole slide images provided by The Cancer Genome Atlas (TCGA) [34]. The BCSS dataset was built on the collaborative effort of pathologists including senior and junior regents, and medical students of pathology by using the Digital Slide Archive, a web-based platform for whole-slide digital pathology images. 151 large ROIs of whole-slide images were annotated by them enabling the generation of accurate deep-learning models for tissue region segmentation.

We obtained 151 large ROIs extracted at 20x magnification from the BCSS dataset. A total of 2,355 H&E images were cropped to 256×256 size and resolution from the 151 large image ROIs. These H&E images were used as the dataset for validating the performance of the proposed SN-FPN on the tumor and stroma region segmentation. To perform a tumor region segmentation, we labeled all the pixels as ‘non-tumor’ if the pixels are not annotated as a tumor. For training SN-FPN, the 2,355 H&E images were randomly divided into three datasets: training, validating, and testing, with 1,413(60%), 417(20%), and 471(20%) respectively.

#### Baseline and metrics

We compare the proposed SN-FPN with the state-of-the-art deep learning-based semantic segmentation models including DeepLabV3Plus [45], UNet++ [39], LinkNet [41], MANet [40], PAN [43], PSPNet [42], and FPN [24], using ResNet50 as an encoder pathway. We use them as the baselines to validate the effectiveness of the proposed model. We used a training epochs of 20, batch size of 8, and learning rate of 0.0001 DeepLabV3Plus is an extended version of the DeepLabV3 model [44] developed by Google Research teams and used for semantic segmentation. DeepLabV3Plus is well-known for capturing multi-scale contextual information by using dilated convolutions integrating with an atrous spatial pyramid pooling [45]. UNet++ is an extended version of the UNet model [38] used for capturing contextual information through the encoder and decoder networks. UNet++ especially uses an adaptive feature selection to dynamically select relevant features at different resolutions. LinkNet is a semantic segmentation model using the encoder and decoder networks. LinkNet uses some link modules for refining high-resolution information by providing an efficient upsampling method. MANet is a multi-scale attention network developed by Tongle et al [40]. MANet uses a self-attention mechanism for integrating local features by capturing contextual dependencies rather than applying the multi-scale feature combination. PAN is a pyramid attention network presented for exploiting and capturing global contextual information in semantic image segmentation [43]. PSPNet is a pyramid scene-parsing network aggregating multi-region context information and providing pyramid pooling models [42]. We use these models for our baselines. We use well-known evaluation metrics such as Intersection over Union (*IoU*), F score (*F* 1), Accuracy (*ACC*), Precision (*PREC*), and Recall (*REC*). *IoU* is defined as (*I* + *ϵ*)*/U* where *I* represents the number of pixels overlapped between the ground truth pixels and the predicted pixels. *ϵ* is set to 1*e* − 7 and added to avoid zero division. *U* represents the encompassed area by both the ground truth pixels and the predicted pixels. *U* is defined as *N*_*g*_ + *N*_*p*_ − *I* + *ϵ* where *N*_*g*_ is the number of the ground truth pixels and *N*_*p*_ the number of predicted pixels. We use a confusion matrix including true positive: TP, false positive: FP, false negative: FN, and true negative: TN, to compute the *F* 1, *ACC, PREC*, and *REC. PREC* is defined as (TP+*ϵ*)/(TP+FP+*ϵ*), *REC* is defined as (TP+*ϵ*)/(TP+FN+*ϵ*), *ACC* is defined as (TP+TN) / (TP+FP+FN+TN), and *F* 1 is defined as ((1 + *β*^2^) * TP + *β*^2^) / ((1 + *β*^2^) * TP + *β*^2^ * FN + FP + *ϵ*) where *β* is set to 1.

#### Results

The performance results on tumor region segmentation comparing the proposed SN-FPN with baseline models are shown in TABLE 1. The results indicate that the SN-FPN outperforms other models in terms of *IoU* (77.40%), *F* 1(83.17%), *ACC*(91.56%), and *REC*(90.19%), and the PAN outperforms other models in terms of *PREC*(88.66%). Since the precision aims to measure the accuracy of the positive predictions indicating the model produces a lower false positive or a higher true positive, it may not be agreeable for other situations where the model requires a lower false negative. Thus, it is necessary to consider the harmonic averages of prediction and recall including both false positive and false negative. To address this consideration we computed the F score in this experiment and the proposed SN-FPN model showed the best result on tumor region segmentation in terms of *F* 1. Moreover, the result of the higher *IoU* on the proposed SN-FPN model indicates that the predicted tumor segmentation regions are more aligned with the ground-truth tumor regions showing the best localization and accurately delineating the target tumor regions. The examples of the performance comparison on tumor region segmentation are shown in Fig. 4.

**TABLE 1.** Performance Results on Tumor Region Segmentation.

| Types | IoU | F1 | ACC | PREC | REC |
| --- | --- | --- | --- | --- | --- |
| DeepV3Plus [45] | 0.7466 | 0.8067 | 0.9116 | 0.8500 | 0.8539 |
| Linknet [41] | 0.7450 | 0.8051 | 0.9095 | 0.8532 | 0.8555 |
| MANet [40] | 0.7370 | 0.7976 | 0.9029 | 0.8109 | 0.8953 |
| PAN [43] | 0.7425 | 0.8023 | 0.9097 | **0.8866** | 0.8220 |
| PSPNet [42] | 0.7383 | 0.7991 | 0.9087 | 0.8388 | 0.8658 |
| Unet++ [39] | 0.7483 | 0.8085 | 0.9154 | 0.8575 | 0.8510 |
| FPN [24] | 0.7564 | 0.8152 | 0.9140 | 0.8403 | 0.8913 |
| SN-FPN | **0.7740** | **0.8317** | **0.9156** | 0.8478 | **0.9019** |

**FIGURE 4.**
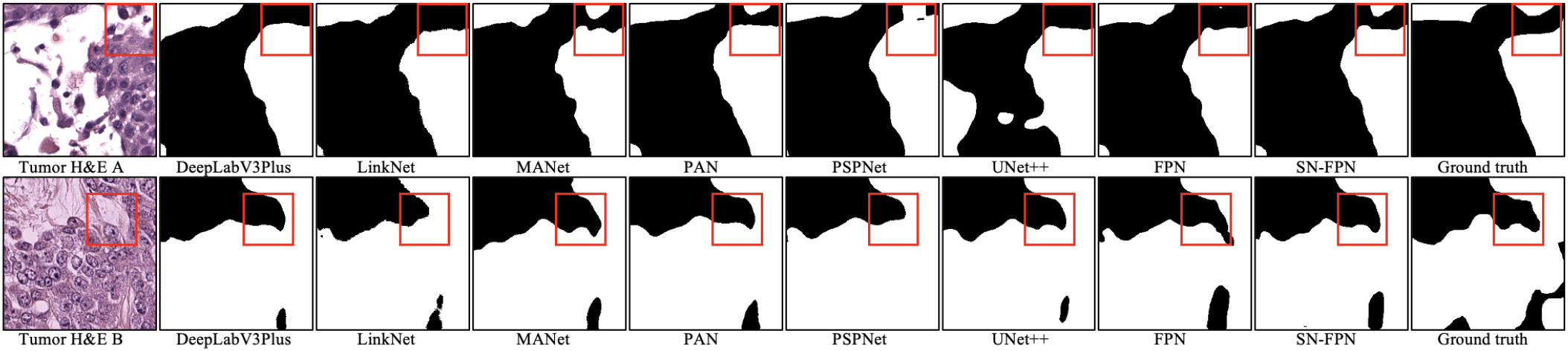
Examples of Tumor Region Predictions using DeepV3Plus, Linknet, MANet, PAN, PSPNet, Unet++, FPN, and SN-FPN.

### B. H&E IMAGE SEGMENTATION FOR STROMA REGIONS

#### Dataset

The 151 large H&E images from the BCSS dataset were used for stroma region segmentation. As we used in the tumor region segmentation, we randomly split the 2,355 H&E images of 256×256 size into 1,413(60%), 417(20%), and 417(20%) H&E images for training, validating, and testing respectively.

#### Baseline and metrics

The baseline models are the same as the models used in the previous section. We compare the models: DeepLabV3Plus [45], UNet++ [39], LinkNet [41], MANet [40], PAN [43], PSPNet [42], and FPN [24] with the proposed SN-FPN model using the same evaluation metrics: *IoU*, *F* 1, *ACC, PREC*, and *REC*.

#### Results

The performance results on stroma region segmentation are shown in TABLE 2. The results indicate that the SN-FPN outperforms other models in terms of *IoU* (54.52%), *F* 1(62.19%), and *PREC*(71.79%). The FPN outperforms other models in terms of *ACC*(86.77%) and the PAN out-performs other models in terms of *REC*(77.93%). Since the accuracy aims to mainly measure the proportion of correctly predicted samples over the total samples, the *ACC* on FPN may not properly provide information about mispredicted samples determined by false positives and false negatives. Moreover, Although *REC* on PAN can provide information about both true positive and false negative samples, it is difficult to mention that PAN outperforms other models because the models should consider the harmonic averages of both prediction and recall. The SN-FPN showed the best *F* 1 score combining precision and recall, indicating that the proposed model outperforms other models on stroma region segmentation using the BCSS dataset. Moreover, the *IoU* score obtained from the SN-FPN represents the proposed model outperforms other models in terms of localization and delineation of the stroma region prediction. The examples of the performance comparison on stroma region segmentation are shown in Fig. 5.

**TABLE 2.** Performance Results on Stroma Region Segmentation.

| Types | IoU | F1 | ACC | PREC | REC |
| --- | --- | --- | --- | --- | --- |
| DeepV3Plus [45] | 0.5404 | 0.6162 | 0.8673 | 0.6884 | 0.7632 |
| Linknet [41] | 0.5372 | 0.6167 | 0.8646 | 0.7108 | 0.7089 |
| MANet [40] | 0.5221 | 0.5982 | 0.8534 | 0.6705 | 0.7577 |
| PAN [43] | 0.5205 | 0.5908 | 0.8659 | 0.6566 | **0.7793** |
| PSPNet [42] | 0.5382 | 0.6173 | 0.8657 | 0.7057 | 0.7193 |
| Unet++ [39] | 0.5443 | 0.6199 | 0.8647 | 0.6997 | 0.7508 |
| FPN [24] | 0.5359 | 0.6097 | **0.8677** | 0.6701 | 0.7788 |
| SN-FPN | **0.5452** | **0.6219** | 0.8533 | **0.7179** | 0.7284 |

**FIGURE 5.**
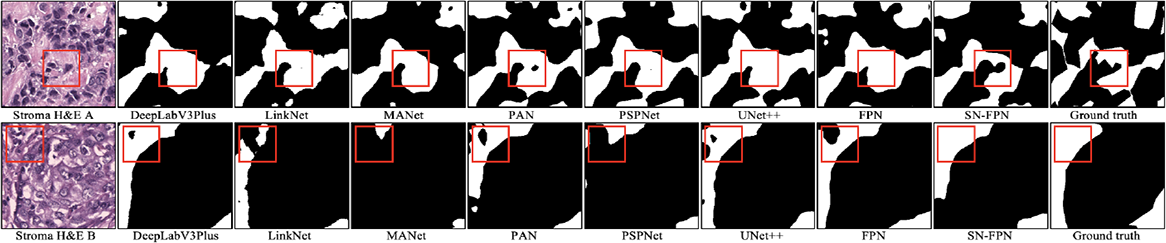
Examples of Stroma Region Predictions using DeepV3Plus, Linknet, MANet, PAN, PSPNet, Unet++, FPN, and SN-FPN.

### C. COMPARISON OF THE ENCODER PATHWAYS ON TUMOR REGION SEGMENTATION

#### Dataset

We used the same dataset of the tumor region segmentation described in Section IV.A. The 2,355 H&E images were used for the comparison of the encoder pathways on tumor region segmentation and divided into three datasets for training 1,413(60%), validating 417(20%), and testing 417(20%) respectively.

#### Baseline and metrics

We conduct experiments on tumor region segmentation using various encoder pathways containing ResNet50 [37], ResNet101 [37], ResNet152 [37], MobileNetv2 [47], EfficientNet-b0, b1, b2, b3, b4, b5, b6, and b7 [46], VGG16 [48], and VGG19 [48]. ResNet101 and ResNet152 are deeper versions of ResNet50 extending layers to 101 and 152 respectively. MobileNet-v2 is an extended version of MobileNet providing an efficient inverted residual block using lightweight depthwise convolutions [47]. EfficientNet is a scaling-emphasized architecture addressing network balance based on different depths, widths, and resolutions thereby leading to better performance [46]. This architecture provides different variants such as b0, b1, b2, b3, b4, b5, b6, and b7. VGG16 and VGG19 are both representative deep convolutional neural network architectures presented by the Visual Geometry Group. VGG16 consists of 16 layers while VGG19 consists of 19 layers [48]. We compare them each other using the tumor region segmentation dataset. We use the same evaluation metrics: *IoU*, *F* 1, *ACC, PREC*, and *REC*.

#### Results

The experiment results on the comparison of the tumor region segmentation using different encoder pathways are shown in TABLE 3. The results on the tumor region segmentation indicate that ResNet50 outperforms other pathways in terms of *IoU* (77.40%), *F* 1(83.17%), and *REC*(90.19%). VGG16 outperforms other encoder pathways in terms of *PREC*(88.22%) and EfficientNet-b6 outperforms other encoder pathways in terms of *ACC*(91.60%). However, the harmonic average *F* 1 of the precision and recall represents that ResNet50 shows better performance on the tumor region segmentation than other pathways in regards to the BCSS dataset.

**TABLE 3.**
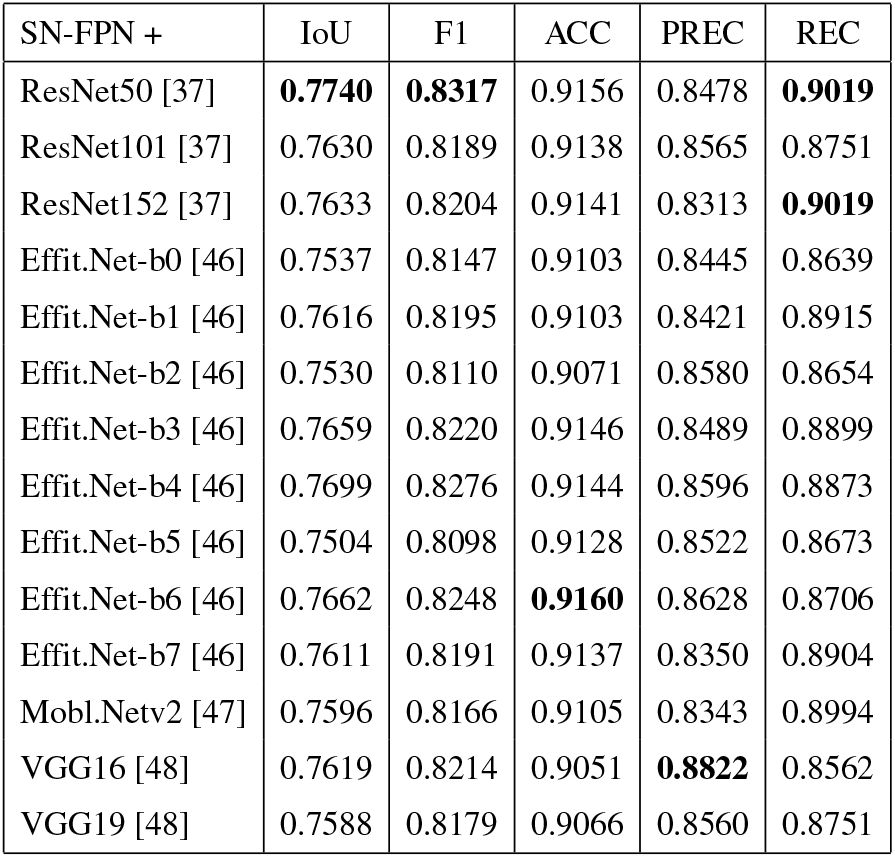
Experiment Results on Tumor Region Segmentation with Different Encoder Pathways.

| SN-FPN + | IoU | F1 | ACC | PREC | REC |
| --- | --- | --- | --- | --- | --- |
| ResNet50 [37] | **0.7740** | **0.8317** | 0.9156 | 0.8478 | **0.9019** |
| ResNet101 [37] | 0.7630 | 0.8189 | 0.9138 | 0.8565 | 0.8751 |
| ResNet152 [37] | 0.7633 | 0.8204 | 0.9141 | 0.8313 | **0.9019** |
| Effit.Net-b0 [46] | 0.7537 | 0.8147 | 0.9103 | 0.8445 | 0.8639 |
| Effit.Net-b1 [46] | 0.7616 | 0.8195 | 0.9103 | 0.8421 | 0.8915 |
| Effit.Net-b2 [46] | 0.7530 | 0.8110 | 0.9071 | 0.8580 | 0.8654 |
| Effit.Net-b3 [46] | 0.7659 | 0.8220 | 0.9146 | 0.8489 | 0.8899 |
| Effit.Net-b4 [46] | 0.7699 | 0.8276 | 0.9144 | 0.8596 | 0.8873 |
| Effit.Net-b5 [46] | 0.7504 | 0.8098 | 0.9128 | 0.8522 | 0.8673 |
| Effit.Net-b6 [46] | 0.7662 | 0.8248 | **0.9160** | 0.8628 | 0.8706 |
| Effit.Net-b7 [46] | 0.7611 | 0.8191 | 0.9137 | 0.8350 | 0.8904 |
| Mobl.Netv2 [47] | 0.7596 | 0.8166 | 0.9105 | 0.8343 | 0.8994 |
| VGG16 [48] | 0.7619 | 0.8214 | 0.9051 | **0.8822** | 0.8562 |
| VGG19 [48] | 0.7588 | 0.8179 | 0.9066 | 0.8560 | 0.8751 |

### D. COMPARISON OF THE ENCODER PATHWAYS ON STROMA REGION SEGMENTATION

#### Dataset

We used the same dataset (2,355 H&E images) of the stroma region segmentation described in Section IV.B. for the comparison of the encoder pathways on stroma region segmentation. The images were divided into three datasets for training 1,413(60%), validating 417(20%), and testing 417(20%) respectively.

#### Baseline and metrics

We use the same encoder pathways: ResNet50 [37], ResNet101 [37], ResNet152 [37], MobileNetv2 [47], EfficientNet-b0, b1, b2, b3, b4, b5, b6, and b7 [46], VGG16 [48], and VGG19 [48] for stroma region segmentaion and the same methods: *IoU*, *F* 1, *ACC, PREC*, and *REC* will be used as the evaluation metrics.

#### Results

The experiment results on the comparison of the stroma region segmentation using different encoder pathways are shown in TABLE 4. The results on the stroma region segmentation indicate that ResNet152 outperforms other pathways in terms of *IoU* (54.95%), *ACC*(87.40%), and *REC*(80.93%) and ResNet50 outperforms other pathways in terms of *F* 1(62.19%). VGG16 shows the best results on *PREC*(76.44%). Assuming that the overall performance is measured by *F* 1 score demonstrating the balance between the precision and the recall, we can determine that ResNet50 shows better performance on the stroma region segmentation than other encoder pathways in regards to the BCSS dataset.

**TABLE 4.** Experiment Results on Stroma Region Segmentation with Different Encoder Pathways.

| SN-FPN + | IoU | F1 | ACC | PREC | REC |
| --- | --- | --- | --- | --- | --- |
| ResNet50 [37] | 0.5452 | **0.6219** | 0.8533 | 0.7179 | 0.7284 |
| ResNet101 [37] | 0.5357 | 0.6109 | 0.8603 | 0.7082 | 0.7281 |
| ResNet152 [37] | **0.5495** | 0.6218 | **0.8740** | 0.6617 | **0.8093** |
| Effit.Net-b0 [46] | 0.5435 | 0.6172 | 0.8654 | 0.6790 | 0.7602 |
| Effit.Net-b1 [46] | 0.5369 | 0.6104 | 0.8674 | 0.6768 | 0.7660 |
| Effit.Net-b2 [46] | 0.5318 | 0.6055 | 0.8553 | 0.6888 | 0.7418 |
| Effit.Net-b3 [46] | 0.5360 | 0.6080 | 0.8688 | 0.6683 | 0.7694 |
| Effit.Net-b4 [46] | 0.5462 | 0.6216 | 0.8659 | 0.6809 | 0.7540 |
| Effit.Net-b5 [46] | 0.5340 | 0.6070 | 0.8667 | 0.6599 | 0.7731 |
| Effit.Net-b6 [46] | 0.5479 | 0.6202 | 0.8665 | 0.6746 | 0.7754 |
| Effit.Net-b7 [46] | 0.5361 | 0.6089 | 0.8674 | 0.6854 | 0.7541 |
| Mobl.Netv2 [47] | 0.5388 | 0.6166 | 0.8712 | 0.6922 | 0.7440 |
| VGG16 [48] | 0.5358 | 0.6155 | 0.8329 | **0.7644** | 0.6877 |
| VGG19 [48] | 0.5306 | 0.6021 | 0.8497 | 0.6737 | 0.7640 |

### E. EXPLAINABLE-AI PERFORMANCE IN PATHOLOGICAL IMAGES

Machine learning models are considered black box models due to their ambiguity in the decision-making process. Explainable AI methods are developed to understand the rationale behind a model decision for prediction. We can also understand whether models are learning key features that might be related to objects or unrelated spurious features of input datasets. Overall, an explainable AI model can help us debug a machine learning model to improve and verify its performance. Explainable AI methods play critical roles in medical images as they can justify the decision provided by machine learning models. Doctors can verify the Machine learning model’s prediction before making crucial patient decisions.

GradCAM [48] is a popular class activation mapping (CAM) method that uses gradients of the last convolution layer to identify essential regions for a given class. Grad-CAM feeds the image to the well-trained deep convolutional network (DCNN) based model to generate the explanation map for an image by using activation functions for the target class. We also use the ScoreCAM [49] method, which uses confidence weights of the activation maps of the last convolution layers instead of unstable gradients to generate explanation maps. Once we create the explainable AI maps of each class for the trained model, we use metrics to compare the performances of explainable AI methods instead of relying on human evaluations. Specifically, we used ROAD [50] and Infidelity [51] scores to compare the performances of GradCAM and ScoreCAM. ROAD method removes the features based on the generated saliency maps to identify the drop in model performance. The higher percentage of accuracy drops indicates a better relationship between the dropped features and the model behavior. The infidelity score also shows how the saliency map indicated region relates to model behavior. A lower score of infidelity indicates higher agreement with the saliency maps indicated region for deep learning model behavior.

#### Dataset

We used 406 and 1007 number of Tumor and Non tumor images to train the Deep learning classifier. We randomly balanced the tumor and non tumor images to train the model. Also, we utilized 333 and 138 number of nontumor and tumor images for validations. Then, we evaluated our explainable AI method and metrics on a separated 341 and 130 number of non-tumor and tumor images.

#### Baseline and metrics

we used a ResNet-50 architecture as a backbone to train the deep learning model classifier for tumor and non-tumor images detection. We used a training epochs of 150, image size 224 *×* 224 *×* 3, batch size of 32

#### Results

We achieved a test accuracy of 92.36% on the testing dataset using the ResNet-50 architecture. Then, we fixed the trained model weight to generate the explanations maps of tumor and non-tumor classes using gradCAM and ScoreCAM methods. To identify the better explainable AI maps, we compare with two explainable AI metrics: ROAD and infidelity. We used ROAD metrics to evaluate the performance of the explainable AI models in Table 5. ScoreCAM performed better than the GradCAM after removing 20% to 90% of the dataset based on explanation maps. Accuracy of the model dropped to 72.82% and 69.64% for the GradCAM and ScoreCAM respectively in Table 5. We used infidelity to compare between GradCAM and ScoreCAM method for tumor detection. We achieved infidelity score of 4.546835771179758*e*^−05^ and 7.94731022324413*e*^−05^ for tumor class using GradCAM and ScoreCAM respectively. We also achieved 1.0336164450563956*e*^−05^ and 9.88344254437834*e*^−05^ for non tumor class using GradCAM and ScoreCAM respectively.

**TABLE 5.** Accuracy of Model After Removing Different Percentages (ROAD Metric) for GradCAM and ScoreCAM Methods on Tumor and non-tumor image Detection.

| % Removed | GradCAM | ScoreCAM |
| --- | --- | --- |
| 0% | 92.36% | 92.36% |
| 10% | 89.81% | 89.81 % |
| 20% | 87.69% | 86.41 % |
| 30% | 85.99% | 81.74 % |
| 40% | 83.44% | 79.83 % |
| 50% | 82.17 % | 78.13 % |
| 70% | 79.41 % | 75.80 % |
| 90% | 72.82 % | 69.64 % |

The ScoreCAM model performed better than the Grad-CAM method in ROAD metrics. If we compare the performance of infidelity-based metrics, we find that both methods performed similarly, and GradCAM is slightly better than the ScoreCAM method.

## V. CONCLUSION

In this paper, we presented a new image segmentation model integrating a feature pyramid network with a self-attention mechanism using three pathways: the encoder pathway, the decoder pathway, and the self-attention nested pathway, and providing accurate tumor and stroma region segmentation on whole slide images. ResNet50 architecture was used for the encoder pathway, while the feature pyramid network was used for the decoder pathway. The self-attention nested pathway generates the attention map from the distribution of attention scores focusing on relevant information of the feature maps. The experiment results show that the proposed SN-FPN model outperforms the state-of-the-art image segmentation models when performing tumor and stroma region segmentation on the BCSS dataset. Moreover, we compared various encoder networks by using the proposed SN-FPN model. The results indicate that the ResNet50 outperforms other encoder networks on F1 score for the tumor and stroma region segmentation.

## ACKNOWLEDGMENT

This research was partially supported by the National Science Foundation under Grant No. 2409704 and Grant No. 2409705.

**SANGHOON LEE** (Member, IEEE) received the B.S. degree in computer science from University of Suwon, South Korea, in 2004 and the M.S. and Ph.D. degree in computer science from Georgia State University, Atlanta, Georgia, in 2014 and 2016 respectively. He is currently an Assistant Professor at Kennesaw State University, Marietta, GA, USA. His research interests include computer vision, medical image processing, machine learning, and deep learning.

**KAZI AMINUL ISLAM** Kazi Aminul Islam received the B.Sc. Degree in Electrical and Electronic Engineering from Khulna University of Engineering and Technology (KUET) in 2010 and master’s degree in Electrical Engineering from Lamar University in 2016. He completed his PhD in Electrical and Computer Engineering from Old Dominion University in 2021. He was a research assistant professor in school of cybersecurity at Old Dominion University between 2021 to 2022. Since 2023, he is working as an assistant professor of Computer Science department at Kennesaw State University. His research includes AI Security and Privacy, cybersecurity, machine learning/deep learning and image processing. He published 24 peer-reviewed journals and conference papers, e.g., IEEE International Conference on Data Mining (ICDM), Remote Sensing of Environment, Remote Sensing.

**SAI CHANDANA KOGANTI** received the B.Tech. degree in Information Technology from Vignan’s Nirula Institute of Technology and Science for Women, Guntur, India in 2023. She is currently pursuing the M.S. degree in Computer Science at Kennesaw State University, Marietta, GA, USA. Her research interests include machine learning, bioinformatics, medical image analysis, computer vision, natural language processing, and web development. She has published papers in conferences like IEEE ICSC and journals on topics such as ensemble learning for medical image segmentation, number plate detection using OpenCV, and web-based learning systems. Ms. Koganti was a Graduate Research Assistant at Kennesaw State University from 2023-present, where she collaborated on machine learning research projects. She has completed certifications from NPTEL, Infosys, Microsoft and Wipro on topics like Java, Python, Data Analytics and Machine Learning.

**SAI RAMYA SRI MAMILLAPALLI** is currently pursuing Masters in Computer Science (MSCS) in Kennesaw State University. She holds a Bachelor’s degree in Information Technology from Vignan’s Nirula Institute of Technology where she gained comprehensive knowledge in Computer Science and related disciplines. Ms. Mamillapalli has authored research paper titled as DYNAMIC WEIGHTED FEATURE SUBSET LOGISTIC REGRESSION MODEL FOR HEART DISEASE PREDICTION published in renowned journal at International Conference on Intelligent Sustainable Systems [ICISS 2023]. She is keen on learning more about Machine Learning, Natural Language Processing and Web Development. She received the certifications from Microsoft, Coursera and NPTEL in C, Python, HTML and in AI. Driven by curiosity for how technology transforms Ms. Mamillapalli is committed to advancing the fields of Machine Learning and open to collaborate on the projects which aim to develop my skills more in the future.

**VARSHINI YAGANTI** is currently pursuing my Master’s degree in Computer Science at Kennesaw State University. Her research journey has been exciting, focusing initially on object detection in images and crop yield forecasting. She is now shifting my focus towards research in biomaterials, a field that fascinates me deeply. Throughout her academic career, She has had the opportunity to work on various research projects. Ms. Yaganti has published her work in reputable journals such as the IEEE (Institute of Electrical and Electronics Engineers) and UGC (University Grants Commission).

**HANNAH VITALOS** Hannah R. Vitalos became a Member (M) of IEEE in 2022 and was born in Annapolis, Maryland, on December 7, 2001. Vitalos received the B.S. degree in computer science from Marshall University, Huntington, WV, USA, in 2024, and is working on the M.S. in computer science from the same institution. She is presently a Research Assistant at Marshall University. In the summers of 2021 and 2022, she was a Software Development Intern at Amazon. In the summer of 2023, she worked as an OSTEM Intern at NASA. Her research interest include bioinformatics and computational biology.

**DREW FK WILLIAMSON** received a B.A. in mathematics from Oberlin College, Oberlin, OH, USA, in 2011 and an M.D. from Case Western Reserve University School of Medicine, Cleveland, OH, USA, in 2018. He is board certified in Anatomic Pathology and Molecular Genetic Pathology and is currently an Assistant Professor in the Department of Pathology and Laboratory Medicine at Emory University School of Medicine in Atlanta, GA, USA. His research focuses on the application of deep learning methods to multimodal pathology data including images, multiomics, and natural language.

